# Dominating Lengthscales of Zebrafish Collective Behaviour

**DOI:** 10.1101/2021.09.01.458490

**Authors:** Yushi Yang, Francesco Turci, Erika Kague, Chrissy L. Hammond, John Russo, C. Patrick Royall

**Affiliations:** H.H. Wills Physics Laboratory, Tyndall Avenue, Bristol, BS8 1TL, UK and Bristol Centre for Functional Nanomaterials, University of Bristol, Bristol, BS8 1TL, UK; H.H. Wills Physics Laboratory, Tyndall Avenue, Bristol, BS8 1TL, UK; Department of Physiology, Pharmacology, and Neuroscience, Medical Sciences, University of Bristol, Bristol, BS8 1TD, UK; Department of Physics, Sapienza University, P.le Aldo Moro 5, 00185 Rome, Italy; Gulliver UMR CNRS 7083, ESPCI Paris, Université PSL, 75005 Paris, France

## Abstract

Collective behaviour in living systems is observed across many scales, from bacteria to insects, to fish shoals. Zebrafish have emerged as a model system amenable to laboratory study. Here we report a three-dimensional study of the collective dynamics of fifty Zebrafish. We observed the emergence of collective behaviour changing between polarised to randomised, upon adaption to new environmental conditions. We quantify the spatial and temporal correlation functions of the fish and identify two length scales, the persistence length and the nearest neighbour distance, that capture the essence of the behavioural changes. The ratio of the two length scales correlates robustly with the polarisation of collective motion that we explain with a reductionist model of self–propelled particles with alignment interactions.

## I. INTRODUCTION

In living systems aggregation occurs at different scales, ranging from bacteria (microns) to insects (centimetres) to fish shoals (tens of kilometres) and with emerging complex patterns [1, 2]. These manifestations of collective behaviour originate from the interactions among the individual agents and between the agents and the environment [3]. Such interactions are often modelled by a combination of deterministic and stochastic contributions, capturing the individual’s variability observed in nature and unknown or uncontrollable variables. The emergence of collective behaviour has been shown to be advantageous for communities [4–6], and identifying commonalities and universal patterns across scales and species is an outstanding challenge [7, 8]. Understanding the relationship between the collective behaviour and animal interactions has potential technological applications, for example to reverse engineer algorithms for the design of intelligent swarming systems [9]. Successful examples include the global optimisation algorithm for the travelling salesman problem inspired by the behaviour of ants, and implementation of the Boids flocking model in schooling of robotic fish [10, 11].

In a reductionist approach, collective behaviour can be modelled with interacting agents representing individuals in living systems. For example, groups of animals may be treated as if they were self-propelled particles with different interacting rules [12, 13]. Examples of using simple agent–based models applied to complex behaviour include describing the curvature of the fish trajectories as a Ornstein–Uhlenbeck process [14], modelling the ordered movement of bird flocks by an Ising spin model [2, 15], mapping of midge swarms onto particulate systems to explain the scale-free velocity correlations [12, 16, 17] and swarming in active colloids [18, 19]. One of the simplest and approaches is the Vicsek model [20], in which the agents only interact via velocity alignment. Despite its simplicity, a dynamical phase transition from polarised flocking to randomised swarming can be identified, providing a basis to describe collective motion in biological systems [21, 22].

The study of collective behaviour in living systems typically has focused on two-dimensional cases for reason of simplicity, making the quantitative characterisation of three-dimensional systems such as flocks of birds of shoal of fish rare. To bridge this gap, Zebrafish *(Danio rerio)* present a wealth of possibilities [23]: zebrafish manifest shoaling behaviour, i.e. they form groups and aggregates, both in nature and in the laboratory; also, it is easy to constrain the fish in controlled environments for long–time observations. Typically, the response of fish to different perturbations, such as food and illumination, can be pursued [23–25]. Furthermore, genetic modification have been very extensively developed for Zebrafish, giving them altered cognitive or physical conditions, and yielding different collective behaviour [26, 27].

However, tracking Zebrafish in three dimensions (3D) has proven difficult [28]. To the best of our knowledge, previous studies on the 3D locomotion of Zebrafish focussed either on the development of the methodology [29, 30], or were limited to very small group sizes (*N* ≤ 5) [28, 31, 32], while ideally one would like to study the 3D behaviour of a statistically significant number of individuals, representative of a typical community. In the field, zebrafish swim in 3D with group sizes ranging from tens to thousands [33].

Here we report on the collective behaviour of a large group (*N* = 50) of wild-type Zebrafish, captured by a custom 3D tracking system. The observed fish shoals present different behaviours, showing different levels of local density and velocity synchronisation. We identify two well-separated time scales (re-orientation time and state-changing time) and two important length scales (persistence length and nearest neighbour distance) for the Zebrafish movement. The time scales indicate the fish group change their collective state gradually and continuously. The spatial scales change significantly as collective behaviour evolves over time, with strong correlations between spatial correlations and shoaling. Finally, we reveal a simple and universal relationship between the global velocity alignment of the shoals (the *polarisation*) and the the ratio between the two length scales (the *reduced persistence length*). We rationalise this finding through the simulation of simple agent-based models, in which an extra inertia term is added to the Vicsek model. Our findings illustrate complex behaviour in Zebrafish shoaling, with couplings between spatial and orientational correlations that could only be revealed through a full three-dimensional analysis.

## II. RESULTS

### A. Experimental Observation

We tracked the movement of zebrafish from multiple angles using three synchronised cameras. We collected data for fish groups with different ages, with young fish (labelled as Y1–Y4) and old fish (labelled as O1–O4). Figure 1(a) schematically illustrates the overall setup of the experiment, where the cameras were mounted above the water to observe the fish in a white tank in the shape of a parabolic dish, enabling 3D tracking [2, 34–36]. With this apparatus, we extract the 3D positions of the centre of each fish at different time points, with the frequency of 15 Hz. We then link these positions into 3D trajectories. Figure 1(b) presents typical 3D trajectories from 50 young zebrafish during a period of 10 seconds, where the fish group changed its moving direction at the wall of the tank. The zebrafish always formed a single coherent group, without splitting into separate clusters during our observations. Movie S1–S3 are examples of the their movements. Figure 1(c) shows the cumulative spatial distribution of the zebrafish in the tank, during a one-hour observation. It is clear from this figure that the fish tend to swim near the central and bottom part of the tank, and that the density distribution is inhomogeneous. The propensity of zebrafish to swim near the wall was our motivation to use a bowl-shaped fish tank shown in(c), so that there are no corners for the fish to aggregate in, compared to a square-shaped container like conventional aquaria.

**FIG. 1.**
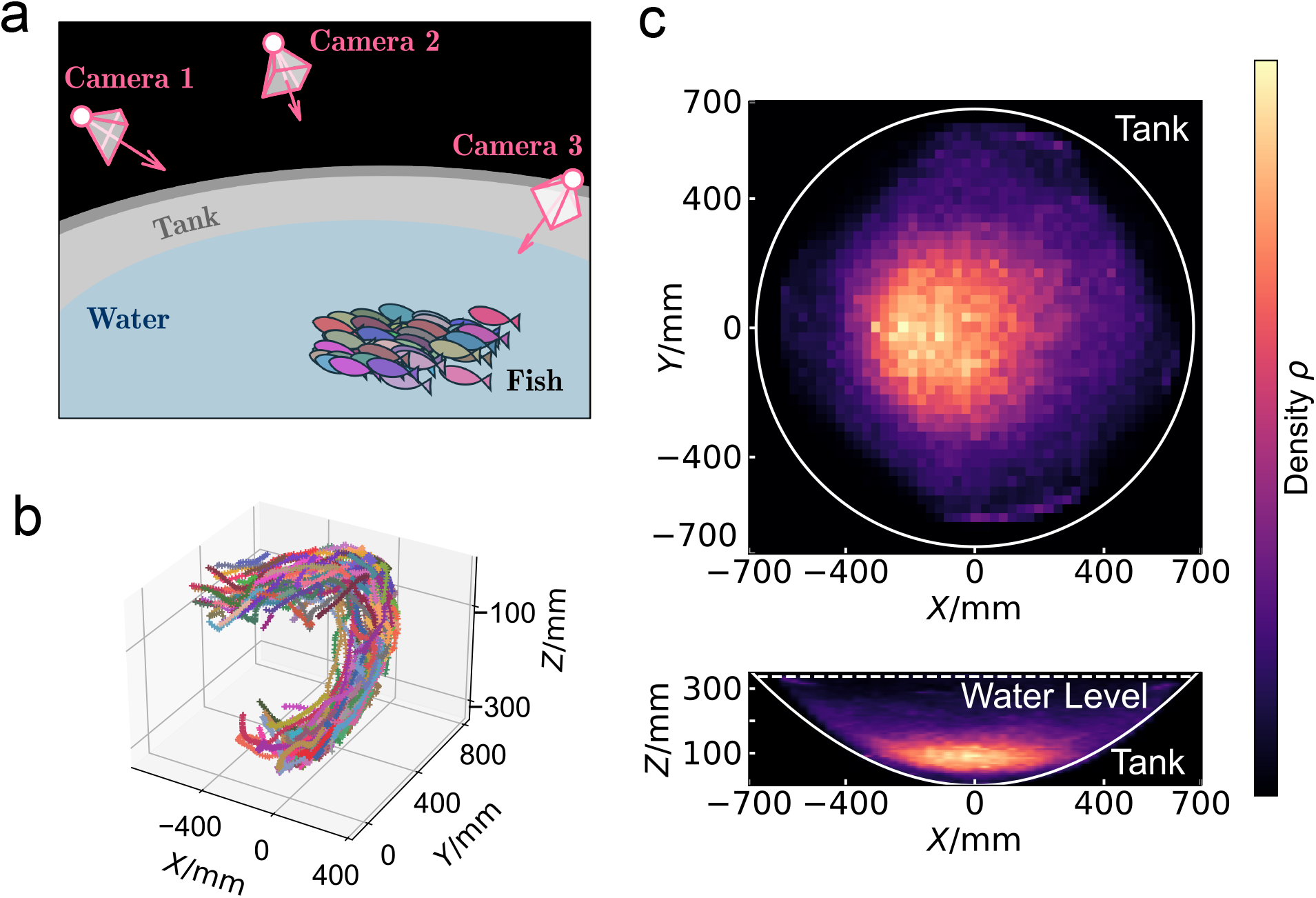
Experimental setup and overall spatial distributions. (a) Schematic illustration of the experimental setup. Zebrafish were transferred into a bowl-shaped tank and three cameras were mounted above the air-water interface to record the trajectories of the fish. (b) 3D trajectories obtained from the synchronised videos of different cameras. These trajectories belong to 50 young Zebrafish (group Y1) in 10 seconds. (c) The spatial distribution of 50 young fish (Y1) during a one-hour observation. Brighter colour indicates higher density. The top panel shows the result in XY plane, obtained from a max-projection of the full 3D distribution. The bottom panel shows a max-projection in XZ plane. The outline of the tank and water-air interface, obtained from 3D measurement, are labelled.

### B. Evolving Collective Behaviour

The 3D tracking yields the positions of the fish, whose discrete time derivative gives the velocities. From these two quantities, we calculate three global descriptors to characterise the behaviour of the fish: the average speed, the polarisation, and the nearest neighbour distance. The average speed is defined as *v*_0_ = 1*/N* Σ|**v**_*i*_| where *i* runs over all the tracked individuals. The polarisation Φ characterises the alignment of the velocities. It is defined as the modulus of the average orientation, written as [20], Φ = 1*/N* |Σ(**v**_*i*_/|**v**_*i*_|)| where *i* runs over all the individuals. Large polarisation (Φ ~ 1) signifies synchronised and ordered movement, while low polarisation indicates decorrelated, random movement. The nearest neighbour distance between the fish centres is defined according to the Euclidean metric, and we focus on is arithmetic mean *d*_1_. We note that while *v*_0_ and Φ are dynamical quantities defined from the velocities *d*_1_ is a static quantity that does not depend on time differences.

We start from the analysis of temporal correlations of these three scalar quantities. Notably, all three exhibit two distinct time scales. Figure 2(a) shows the auto–correlation functions (ACF) of *v*_0_, Φ and *d*_1_ averaged over the group of 50 young fish, calculated from a one hour observation. The ACFs present two decays and one intermediate plateau. We identify the first decay with the short time (~1s) reorientation time, characteristic of Zebrafish behaviour. This can be shown through the analysis of the autocorrelation of the orientations Fig. 2(b), which are characterised by an exponential decay with relaxation time 〈*τ*〉 also close to ~1s, which is compatible with the previously reported turning rate timescale (~0.7s) [37].

**FIG. 2.**
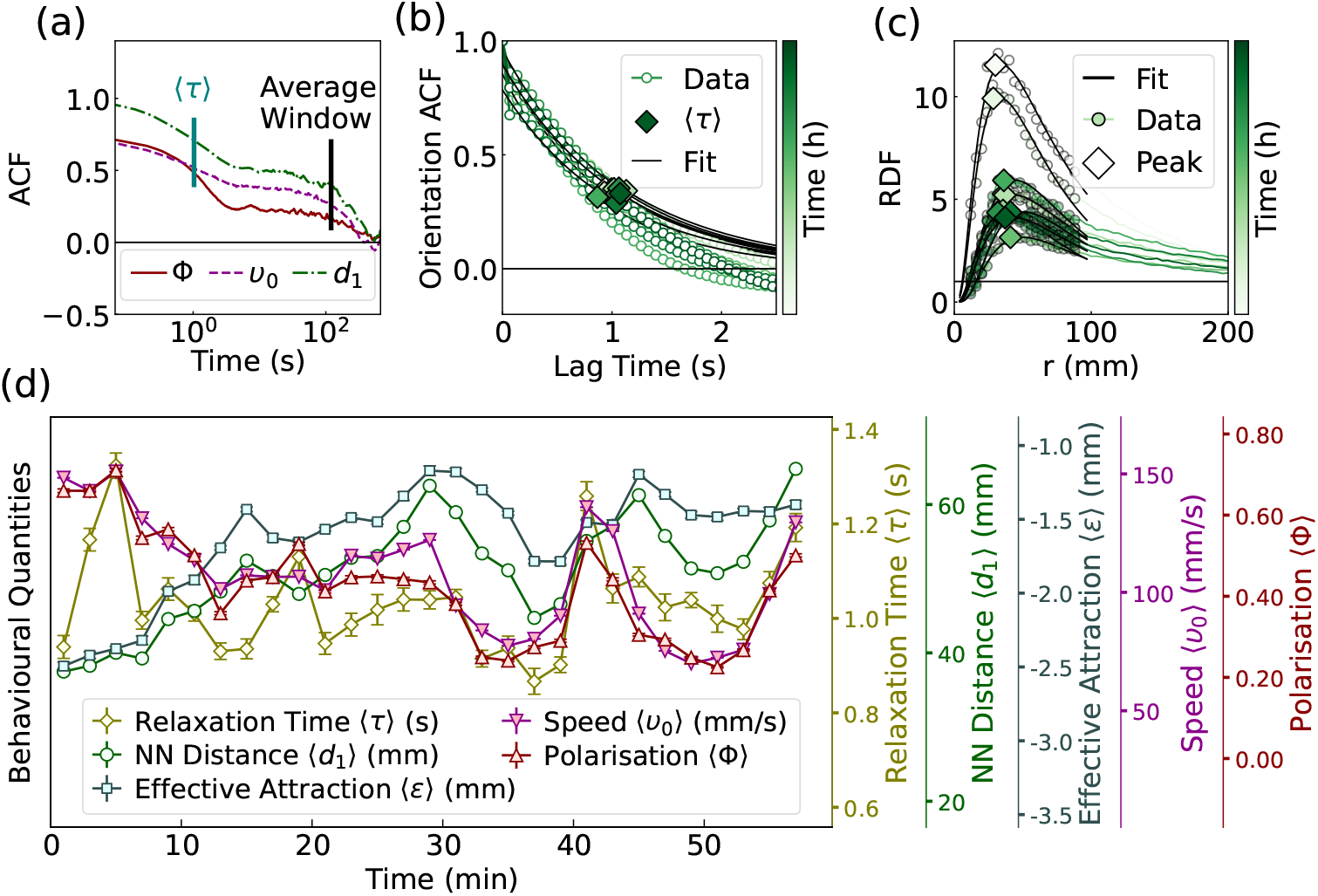
The behavioural quantities of 50 young Zebrafish (group Y1). (a) The auto– correlation function of the polarisation and average speed of the fish group. (b) The auto– correlation function of the orientations of fish. (c) Sequence of radial distribution functions with increasing time: at early times (top curves) the fish are clustered together so that the peak is large; at later times (bottom curves) the local density decreases and so does the peak height. (d) The time evolution of the averaged behavioural quantities for 50 young fish. Each point corresponds to the average value in 2 minutes. The error bars illustrate the standard error values.

The plateau and subsequent decay of the ACF of the scalar quantities *v*_0_, Φ and *d*_1_, with the time scale of ~120 seconds, represent complete decorrelation from the initial state, indicating that the shoal properties change significantly on this much longer timescale. Therefore, we employ time-windows of 120 seconds to average the time evolution of of *v*_0_, Φ and *d*_1_, to characterise the states of the fish groups with moving averages 〈*v*_0_〉(t), 〈Φ〉(t) and 〈*d*_1_〉(t).

To characterise the degree of spatial correlation of the fish, we focus on the fish centre of mass and calculate their radial distribution function (RDF), see Fig. 2(c), which quantifies the amount of pair (fish-fish) correlations and it is commonly employed in the characterisation fo disordered systems ranging from gas to liquids, from plasma to planetary nebulae[38]. Details on the RDF can be found in the supplementary information (SI). All the RDFs exhibit one peak at a short separation, indicating the most likely short-distance separation between fish. The peak height is a measure of the cohesion of the fish group. Inspired by liquid state theory [38], we take the negative logarithm of the peak height to characterise what we call as the “effective attraction” among the fish, noted as 〈*ϵ*〉. While *d*_1_ quantifies a characteristic lengthscale in the macroscopic collective state, *ϵ* quantifies the fish propensity to remain neighbours. In Fig. 2 we see that *d*_1_ and *ϵ* are strongly correlated, confirming that *d*_1_ is also a measure of the cohesion of the collective states. We term 〈*v*_0_〉, 〈Φ〉, 〈*τ*〉, 〈*d*_1_〉, and 〈*ϵ*〉 “behavioural quantities”, and the brackets indicate the moving average.

Figure 2(d) illustrates the time-evolution of all the behavioural quantities, calculated from the movement of 50 young fish (group Y1) ten minutes after they were extracted from a husbandry aquarium and introduced into the observation tank. Over time, the behavioural quantities drift, indicating that a steady state cannot be defined over the timescale of 1 hour. This result is generic, as the separated time scales and changing states were obtained from repeated experiments on the fish group (Y2–Y4), and also from different groups of older Zebrafish (O1–O4).

### C. Shoaling State Described by Two Length Scales

To describe the space of possible collective macroscopic states we employ two dimensioned lengths, the nearest neighbour distance 〈*d*_1_〉 defined above and a second scale characterising the typical distance that a single fish covers without reorientation, the persistence length 〈*l_p_*〉. Tis is defined as the product of the speed and the orientational relaxation time 〈*l_p_*〉 = 〈*v*_0_〉〈*τ*〉.

The resulting 〈*l_p_*〉 and 〈*d*_1_〉 diagram is illustrated in Fig. 3(a). As we move across the diagram, the degree of alignment of the fish motion – the polarisation – also changes, indicating that changes in the local density (as measured by 〈*d*_1_〉) and in the pattern and velocity of motion (as measured by 〈*l_p_*〉) are reflected in the polar order of the shoals. For high 〈*l_p_*〉 and low 〈*d*_1_〉, the movements of the fish are cohesive and ordered (Movie S1). For the fish states with a low 〈*d*_1_〉 and low 〈*l_p_*〉, the movements are cohesive but disordered (Movie S3). For fish states with high 〈*d*_1_〉 and low 〈*l_p_*〉, the fish are spatially separated with disordered movements (Movie S2). Cohesive but dilute states are never observed. We also note that there is a systematic difference between young (Y) and old (O) fish groups, with the former characterised by larger persistence lengths, neighbour distances and polarisations while the latter are clustered in a narrower range of persistence lengths with more disorder.

**FIG. 3.**
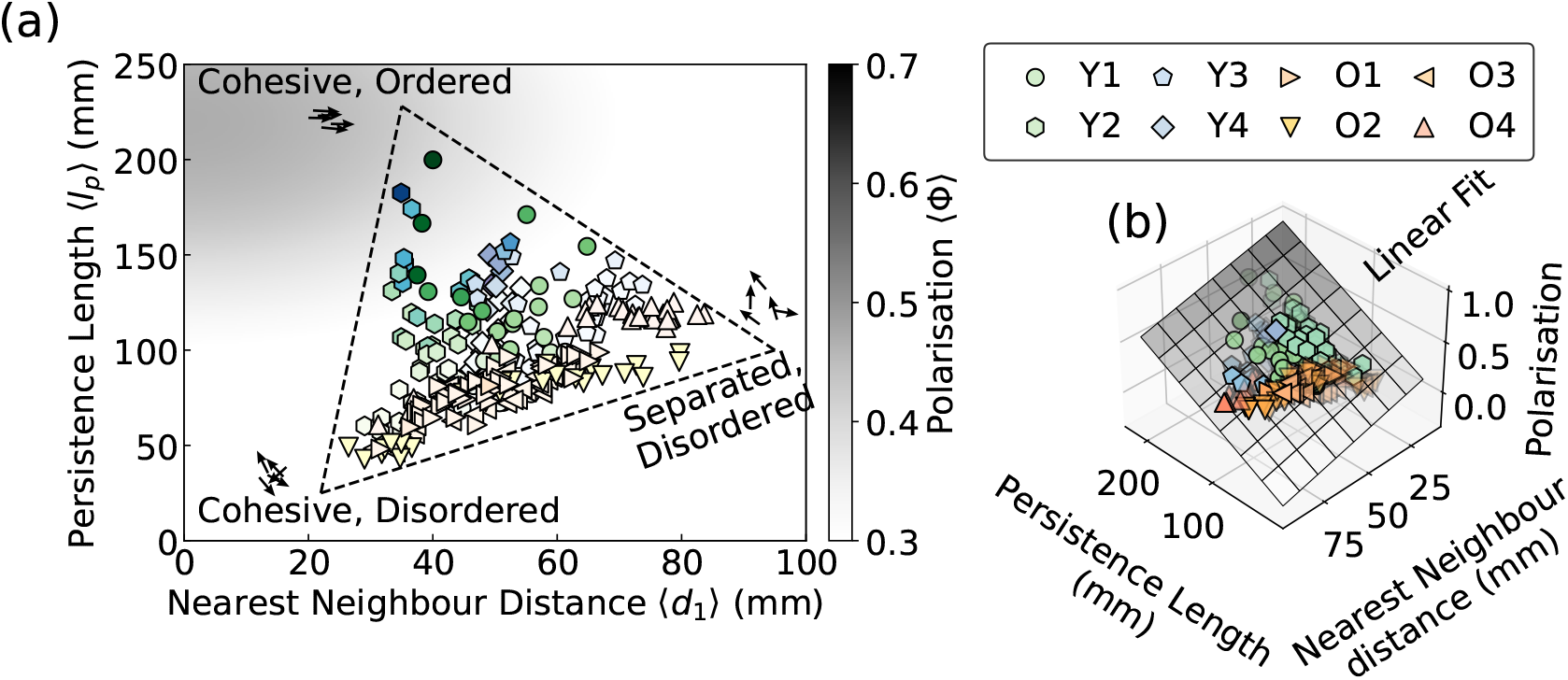
The states of Zebrafish characterised by two length scales. (a) The states of the fish represented by the nearest neighbour distance and the persistence length. The brightness of the markers corresponds to the value of the polarisation. Each scatter–point corresponds to a time-average of 2 minutes. Different shapes indicate different fish groups from different experiments. (b) A multilinear regression model fitting the relationship between the polarisation and the two length scales, indicating the polarisation increase with the increase of persistence length, and the decrease of the nearest neighbour distance. The model is rendered as a 2D plane, whose darkness indicate the value of polarisation.

The simplest model to capture the relationship between polarisation and the two length-scales is a multilinear regression. This yields 〈Φ〉 = 0.039 〈*l_p_*〉 − 0.05 〈*d*_1_〉 + 0.147, with a goodness of fit value (*R*^2^ = 0.73). This strong simplification suggests that most of the fish macroscopic states reside on a planar manifold in the Φ–*d*_1_–*l_p_* space, illustrated in Fig. 3 (b). The value of 〈Φ〉 increases with the increase of 〈*l_p_*〉, and the decrease of 〈*d*_1_〉. Such relationship is reminiscent of results from the agent-based Vicsek model, where the polarisation of self–propelled particles is determined by the density 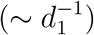 and orientational noise 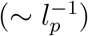 [20, 39]. In addition, the relationship between the polarisation and the local density suggests a metric based interaction rule, rather than the topological counterpart [40]. In other words, the fish tend to align with nearby neighbours, rather than a fixed number of neighbours. A similar relationship between the polarisation and density was also found for jackdaw flocks while responding to predators [41].

A further simplification to describe quantitatively the data can be obtained employing the ratio between the persistence length and the nearest neighbour distance offers a simplified description of the polarisation of the fish groups. Here we introduce the *reduced* persistence length *κ* = 〈*l_p_*〉/〈*d*_1_〉. The value of *κ* exhibits a consistent relationship with the polarisation for all the fish groups, as shown in Fig. 4 (a). All the experimental data points collapse onto a single curve, especially for the younger fish groups (Y1–Y4) which have a much wider dynamic range than the older groups. Notably, the young fish always transform from ordered states with high *κ* value to disordered states with low *κ* value, possibly because of the adaption to the observation tank.

**FIG. 4.**
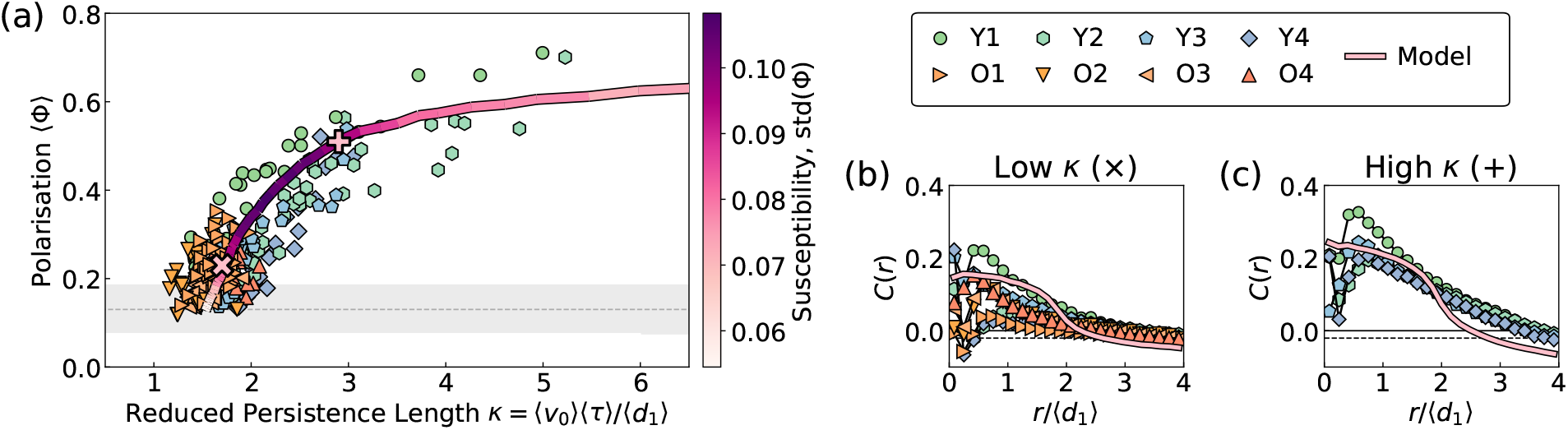
Single–parameter description of the Zebrafish system. (a) The average polarisation 〈Φ〉 as a function of the reduced persistence length *κ*, where most data points collapse, and agree with the simulation result of the inertial Vicsek model. The dashed line and grayscale zone represent the expected average value and standard deviation of 〈Φ〉 for the uniform random sampling of vectors on the unit sphere. (b) The velocity correlation function of the fish and the model in the low *κ* states, highlighted in (a) with a × symbol. (c) The velocity correlation function of the fish and the model in the high *κ* states, highlighted in (a) with a + symbol.

To understand this relationship, we consider the fish motion as a sequence of persistent paths interrupted by reorientations. In a simplified picture, the new swimming direction at a reorientation event is determined by an effective local alignment interaction that depends on the neighbourhood, and notably on the nearest neighbour distance *d*_1_. The fish states with larger value of *κ* correspond to situations where each individual fish interacts with more neighbours on average, between successive reorientations. The increased neighbour number leads to a more ordered collective behaviour, so that the values of *κ* and Φ are positively correlated as shown in Fig. 4(a).

The time-averaged spatial correlation of the velocity fluctuation supported our picture of the local alignment interaction between the fish. Such a correlation function is defined as,

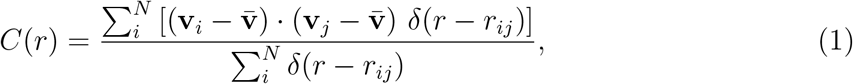

where **v**_*i*_ is the velocity of fish *i*, 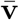 is the average velocity in one frame, *r_ij_* is the distance between two particles, and *δ* is the Dirac delta function. This function is widely used to characterise the average alignment of velocity fluctuations of moving animals, at different distances [2, 42, 43]. Figures 4(b) and (c) show the correlation functions for different fish groups with low and high *κ* values, respectively. The distances are rescaled by the different 〈*d*_1_〉 values of each fish group. For both conditions, the correlation curve collapses beyond one 〈*d*_1_〉, and peaks around the value of 〈*d*_1_〉, supporting our assumption that 〈*d*_1_〉 is the length scale for the fish–fish interactions.

### D. Vicsek Model Rationalisation of the Experiments

The relationship between *κ* and Φ, presented in Fig. 4(a), can be easily compared with simulations as both *κ* and Φ are dimensionless quantities. Here we explore this through simulations proposing a new modification of the original Vicsek model [20]. The Vicsek model treats the fish as point-like agents with an associated velocity vector of constant speed *v*_0_. During the movement, the fish adjust their orientations to align with the neighbours’ average moving direction. In order to take into account of memory effects in a simple fashion, we add an inertia term into the Vicsek model, so that each agent partially retain their velocities after the update, with the following rule

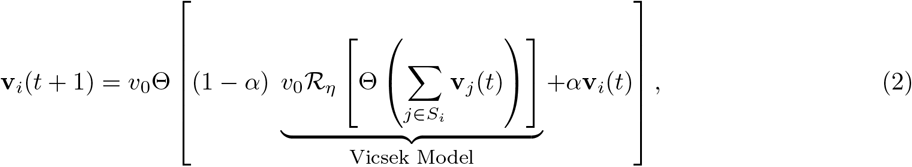

where 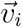 is the velocity or the *i*th fish, and the updated velocity of fish *i* is a linear mixture of its previous velocity and a Vicsek term. The parameter *α* characterises the proportion of the non-updated velocity, *i.e.* the inertia. This model is reduced to the Vicsek model by setting *α* to 0. If *α* = 1, these agents will have constant ballistic movements without any interaction. For the Vicsek term, *S_i_* is the set of the neighbours of fish *i*, and the Θ is a normalising function. The operator 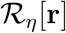 randomly rotates the vector **r** into a new direction, which is drawn uniformly from a cap on the unit sphere. The cap is centred around **r** with an area of 4*πη*. The value of *η* determines the degree of stochasticity of the system. Our model is thus an inertial Vicsek model in three dimensions with vectorial noise.

We set the units of the interaction range *ξ* and time *dt* and fix the number density to *ρ* = 1*ξ*^−3^ and speed *v* = 0.1*ξ/dt*. We then proceed with varying the two parameters *α* and *η* to match the data. In particular we measure the average persistence length 〈*k*〉 and polarisation 〈Φ〉 and find that for *α* = 0.63 we can fit the data only through the variation of the noise strength *η* More details of the simulation are available in the SI). For *η* ~ 1, the movement of each agent is completely randomised, reproducing the low *κ* (or Φ) states of the fish. For the case of *η* ~ 0.65 the movements of the agents are ordered (Φ ~ 0.64) and mimic the states of fish with high *κ*. This is consistent with the fact that in the ordinary Vicsek model the persistence length scales as *ℓ* ~ *v*_0_/*η*^2^. The good fit of the simulation result suggests the fish–fish alignment interaction dominates their behaviour, and the fish can change their states by changing the rotational noise (*η*).

We emphasise that the inertial Vicsek model is a crude approximation, as the only interaction of the model is velocity alignment. Without the attractive/repulsive interactions and other details, the inertial Vicsek model does not reproduce the velocity correlation function of the fish, as illustrated in Fig. 4(b) and(c), suggesting that more sophisticated models with effective pairwise and higher order interactions may be developed in the future. Nevertheless, the model qualitatively reproduces the fact that the velocity correlation is stronger in the high *κ* states.

## III. DISCUSSION AND CONCLUSION

Our results confirm some previous observations and open novel research directions. The young fish appear to adapt to a new environment with the reduction of the effective attraction and speed (Fig. 2). Such behaviour is consistent with previous observations of dense groups of fish dispersing over 2-3 hours [44]. At the same time, it was reported by Miller and Gerlai that the habituation time has no influence on the Zebrafish group density [45]. We speculate that this difference emerges from the way the statistics were performed. Typically, Miller and Gerlai’s results were averaged over 8 different small fish groups (N=16), and it is possible that the noise among different groups obfuscates the time dependence features here highlighted. Despite the different claims, our result matched Miller and Gerlai’s result quantitatively (Fig. S5).

The macroscopic state polarisation of the fish groups decreases during the adaption process for the young fish. This “schooling to shoaling” phenomenon has been observed previously in a quasi 2 dimensional environment [46]. Our results suggest that this behaviour is present also in a fully three-dimensional context and that the change from schooling (polarised motion) to shoaling (unpolarised motion) is related to an increasingly disordered or uncorrelated behaviour, corresponding to the increase in the noise term *η* in the Vicsek model.

It is been speculated that all the biological systems were posed near the critical state, to enjoy the maximum response to the environmental stimuli [47]. Here the inertial Vicsek model offered a supporting evidence. The fluctuation of the polarisation, the *susceptibility*, took a maximum value at moderate reduced persistence value *κ* ~ 2, as illustrated Fig. 4 (a). And the fish states were clustered around such region, where the fish can switch between the disordered behaviour and ordered behaviour swiftly. Such disordered but critical behaviour was also observed for the midges in the urban parks of Rome[17].

In conclusion, our work presents a quantitative study of the spatial and temporal correlations manifested by a large group of Zebrafish. In our fully 3D characterisation, we have shown that there is a timescale separation between rapid reorientations at short times and the formation of a dynamical state with characteristic spatial correlations at longer times. Such spatial correlations evolve continuously and no steady state is observed in the time window of one hour. Our analysis shows that the continuously changing collective macroscopic states of the fish can be described quantitatively by the persistence length and nearest neighbour distance. The ratio of these length scales presents a characteristic correlation with the polarisation of the fish group. This simple relation is supported by an elementary agent based model in the class of the Vicsek models for collective behaviour.

Our analysis also opens multiple questions: the true nature of the interactions and how these are linked to the sensory and vision capabilities of the fish is open to debate; also, the reason for the change of the fish states remained unexplored, with the possibility of the fish learning over time about the experimental conditions. Our work shows that Zebrafish provides a viable model system for the study of animal collective behaviour where such questions can be investigated in a quantitative manner.

A further intriguing possibility is to link the methodology that we develop here, with genetic modifications to Zebrafish, for example with behavioural phenomena such as autism [26] or physical alterations such as the stiffened bone and cartilage [48].

## IV. METHODS

### A. Data and Code Availability

The code for tracking the fish, including the 2D feature detection, 3D locating, trajectory linking, and correlation calculation, is available free and open–source (link to GitHub). The simulation code is also available on GitHub (link to repository). The dataset for generating Fig. 2, 3, and 4 are available as supplementary information (Dataset S1), as well as some pedagogical code snippets (Code S1).

### B. Zebrafish Husbandry

Wildtype Zebrafish were kept in aquarium tanks with a fish density of about 10 fish / L. The fish were fed with commercial flake fish food (Tetra Min). The temperature of the water was maintained at 25°C and the pH ≈ 7. They were fed three times a day and experience natural day to night circles, with a natural environment where the bottom of the tank is covered with soil, water plant, and decorations as standard conditions [49]. Our young group (Y) were adults between 4-6 months post-fertilisation, while our old group (O) were aged between 1-1.5 year. The standard body lengths of these fish were are available in the SI. All the fish were bred at the fish facility of the University of Bristol. The experiments were approved by the local ethics committee (University of Bristol Animal Welfare and Ethical Review Body, AWERB) and given a UIN (university ethical approval identifier).

### C. Apparatus

The movement of the Zebrafish were filmed in a separate bowl-shaped tank, which is immersed in a larger water tank of 1.4 m diameter. The radius *r* increasing with the height *z* following *z* = 0.2*r*^2^. The 3D geometry of the tank is measured experimentally by drawing markers on the surface of the tank, and 3D re-construct the positions of the markers. Outside the tank but inside the outer tank, heaters and filters were used to maintain the temperature and quality of the water. The videos of Zebrafish were recorded with three synchronised cameras (Basler acA2040 um), pointing towards the tank. A photo of the setup, and more details are available in the SI.

### D. Measurement and Analysis

Fifty Zebrafish were randomly collected from their living tank, moved to a temporary container, then transferred to the film tank. The filming started about 10 minutes after fish were transferred. The individual fish in each 2D images were located by our custom script and we calculated the 3D positions of each fish following conventional computer vision method [50, 51].

The 3D positions of the fish were linked into trajectories [52, 53]. Such linking process yielded the positions and velocities of different fish in different frames. We segmented the experimental data into different sections of 120 seconds, and treat each section as a steady state, where the time averaged behavioural quantities were calculated. More details on the tracking and analysis are available in the SI.

## ACKNOWLEDGMENTS

CLH and EK were funded by Versus Arthritis grants 21937 and 21121. YY were funded by the China Scholarship Council (CSC). We thank James G. Puckett, Christos C. Ioannou, and James Herbert for stimulating conversations.

